# Dietary zinc restriction induces nociceptive pain with reduced inflammation in mice

**DOI:** 10.1101/2020.11.21.392548

**Authors:** Cleverton Kleiton Freitas de Lima, Tháyna Sisnande, Rafaela Vieira da Silva, Vanessa Domitilla de Castro Soares da Silva, Julio Jablonski do Amaral, Soraya de Mendonça Ochs, Bruna Lima Roedel dos Santos, Ana Luisa Palhares de Miranda, Luis Mauricio T. R. Lima

**Author notes:** Both authors share senior authorship (Ana Luisa Palhares de Miranda, Luis Mauricio T. R. Lima). To whom correspondence should be addressed: Luis Maurício T. R. Lima – Faculty of Pharmacy, Federal University of Rio de Janeiro – UFRJ, CCS, Bss24, Ilha do Fundão, 21941-902, Rio de Janeiro, RJ, Brazil. Twitter: @MauTrambaioli, Ana Luisa P. Miranda – Faculty of Pharmacy, Federal University of Rio de Janeiro – UFRJ, CCS, Bss12, Ilha do Fundão, 21941-902, Rio de Janeiro, RJ, Brazil.

## Abstract

Zinc (Zn) is an essential micronutrient involved in a large diversity of cellular metabolism, included in the physiology of nervous system and pain modulation. There is little evidence for the role of Zn nutritional alternations to the onset and progression of neuropathic and inflammatory pain. We investigate the effects of a zinc restricted diet on the development of pain. Weaned mice were submitted to different diets: AIN-93 (38mg/kg of Zn) and Zn-deficient (AIN-93 with 11mg/kg of Zn), during four weeks. Mechanical allodynia was measured weekly using Von Frey hairs. Plantar assays for cold and heat allodynia, formalin-induced nociception and carrageenan-induced mechanical allodynia were performed at the 4^th^ week. Plasma, DRG and livers samples were obtained for biochemical and metabolomics analysis. Zn deficient diet completely changed mice sensitivity pattern, inducing an intense allodynia evoked by mechanical, cold and heat stimulus since weaning and during four weeks. Showed also an increased sensitivity of neurogenic phase of formalin test but the inflammatory pain behavior was drastically reduced. Zn restriction increased the ATF-3 and SOD-1 levels at DRG and reduced that of GFAP, leading an increase of neuronal activation and oxidative stress, and reduced neuroimmune activity. Plasma TNF was also reduced and metabolomics analyses suggest a downregulation of lipid metabolism of arachidonic acid, reinforcing the impact of Zn restriction to the inflammatory response. Reduction of Zn intake interferes in pain circuits, reducing inflammatory pain, however enhancing nociceptive pain. Accordingly, Zn imbalance could be predisposing factor for NP development. Therefore, dietary zinc supplementation and its monitoring present clinical relevance.

## 1. Introduction

Zinc is a trace element largely present on human body, being the second most abundant elements within human cells (Fraga, 2005). It is an essential micronutrient involved in a large diversity of cellular metabolism, acting as a catalytic, co-catalytic or structural element of proteins and enzymes. DNA synthesis, cell division and immune response are some examples of biological processes that depends on zinc (Andreini et al., 2006).

The homeostasis of zinc is a vital process to the cell functions and physiology maintaining. Zinc is present in a wide variety of foods as nuts, meats and milk. Exogenous zinc is absorbed primarily in gastrointestinal system, followed by uptake in enterocytes and subsequent distribution through tissues. Endogenous levels of zinc are regulated by a refined system that involves two different families of zinc transporters proteins: ZnTs and ZIPs. The firsts are generally involved in sequestrate this ion into intracellular region or export zinc out of the cell when concentration is high, while ZIP proteins are responsible to bring zinc into cell or release when zinc stores are low (depleted conditions) (Tepaamorndech et al., 2014).

The recommended daily ingestion of zinc to adults should be about 10 mg (Institute of Medicine, 2005). However, about 17% of the world’s population consumes inadequate amounts of zinc (Wessells and Brown, 2012). Delayed wound healing, male hypogonadism, mental lethargy, cell mediated immune dysfunctions and growth retardation are some of the several disorders associated with zinc deficiency (King, 2011; Prasad, 1991).

It has been demonstrated that zinc deficiency results in reduced circulating IGF-1 (insulin-like growth factor 1), which regulates cell cycle and division (Cossack, 1991; Ninh et al., 1995). Furthermore, zinc acts also as a cofactor of tyrosine kinases, enzymes responsible to activate IGF-1 receptor (Wilson et al., 2012). A recent study of our group revealed that zinc restriction produces selective degeneration of the endocrine pancreas in mice (SISNANDE et al., 2020). In the central nervous system (CNS), zinc is an essential micronutrient for the development of memory and learning processes, as well as mediating synaptic plasticity (Hagmeyer et al., 2015). Zinc is expressed in presynaptic terminals of glutamatergic synapses, being co-stored with glutamate in specific vesicles that are mainly localized in cerebral cortex and in the limbic system. In these regions, high densities of a zinc-sensitive metabotropic receptor (ZnR) were detected, as well as ZnT and other proteins considered “zinc buffers” such as zinc import (ZIP) and metallothionein (MT) (Hara et al., 2017). Peripheral nervous system also counts with a variety of processes modulated by zinc. Zinc is present in thoracic and lumbar dorsal root ganglia (DRG), being also accumulated in sensory fibers (Velazquez et al., 1999).

It’s well known that zinc plays important roles in pain modulation. Liu et al (1999) showed that zinc ions delivered by intrathecal, local or systemic injection is able to reduce thermal hyperalgesia in rats with sciatic nerve injury (Liu et al., 1999). Nanomolar concentrations of zinc were able to block NMDA glutamate receptor, reducing neuronal activity in the spinal cord and providing an analgesic response in inflammatory and neuropathic pain models (Nozaki et al., 2011). A recent study also demonstrated that local delivery of zinc inhibits mechanical hypersensitivity evoked by paclitaxel-induced neuropathy, confirming the function of this micronutrient in different kinds of pain (Luo et al., 2017).

Beyond the direct modulation of exogenous zinc in nociceptive models, endogenous zinc levels also seem to be important to pain response. The reduction of zinc levels promoted by chelators agents presented a pro-nociceptive effects in different experimental models (Larson and Kitto, 1997)(Matsunami et al., 2011). In addition, some clinical studies also correlates painful conditions as fibromyalgia and myofascial pain with low levels of plasmatic zinc (Sendur et al., 2008) (Barros-Neto et al., 2016). Acute painful episodes in children with sickle cell anemia was also associated with low serum zinc levels (AA Kudirat, UA Shehu, E Kolade, 2019).

Despite of the growing knowledge about the role of zinc in the CNS and the influence of its dyshomeostasis on the development of degenerative diseases (Szewczyk, 2013)(Wojtunik-Kulesza et al., 2019), the scientific literature presents little evidence for the role of nutritional alternations of this metal to nociceptive disorders, such as neuropathic and inflammatory pain.

Considering the importance of zinc homeostasis for the nervous and immune system functioning, knowing that the imbalance of these systems pivotal to the pathogenesis of different kinds of pain, the present study aimed to investigate the impact of zinc diet restriction in the onset of nociceptive disorders in mice since weaning.

## 2. Material and Methods

### 2.1. Material

The diets were prepared by Rhoster^®^ Ind. Com. LTDA (São Paulo). The Zn content in the diets were determined by mineral analysis (Laboratório de Ciência e Tecnologia, São Paulo, Brazil), revealing a zinc content of 38 mg/kg in the control diet and 11 mg/kg in the intervention diet). All other reagents were from analytical grade and used as received.

### 2.2. Experimental groups

Studies were performed using 21 days old male Swiss mice. Animals were housed at UFRJ’s Faculty of Pharmacy vivarium, with free access to food and water in a 12-hour dark-light cycle. All experimental protocols were approved by the Ethical Animal Care Committee of Federal University of Rio de Janeiro (CEUA-UFRJ) under the number CC-UFRJ-011/2015.

The animals (n=6-8 animals/group), immediately after weaning, were divided into two groups that received regular or zinc deficient diets, and maintained with the assigned diets throughout the experiment. Control group was exclusively fed with the AIN-93 modified chow (with casein replaced by hen egg white solids; Salto’s LTDA), formulated with either regular mineral mix (AIN-93G-MX) or zinc-deficient mineral mix formulated without added zinc salts (AIN-93G-MX-ZnDef) as previously reported (Sisnande et al., 2020).

The body mass and mechanical allodynia were measured weekly. Tibial growth, thermal heat hyperalgesia and thermal cold allodynia were performed on the 4^th^ week. Formalin and carrageenan evoked hyperalgesia tests were performed in independent groups.

### 2.3. Behavior hypersensitivity tests

#### 2.3.1. Mechanical allodynia

The mechanical threshold evaluation was performed as described by (Decosterd and Woolf, 2000). Briefly, animals were acclimated and then submitted to sequential mechanical stimulus ranging from 0.008 g to 2.0 g by *von Frey* filaments. The stimulus was applied under the left hind paw and each filament was presented 5 times to the paw, respecting a 60s gap between then. The mechanical threshold was defined as the last filament that obtained the highest number of paw withdrawal responses. Animals with a paw withdrawal threshold (PWT) higher than 2.0 g were excluded from the experimental groups (outliers). Mechanical sensitivity was measured weekly.

#### 2.3.2. Thermal heat hypernociception

To evaluate the thermal heat *hypernociception*, animals were placed in a plantar thermal testing machine that consists in a contention chamber with a glass floor. After 30 min of acclimation, a mobile heating source (high intensity light) was positioned under the glass floor with the light beam straight to center of the hind paw. Heating stimulus was interrupted with paw withdrawal (nociceptive response) and the time between the light emission and paw withdrawal was considered as the latency time (in seconds). The light beam generated a rate of 1°C/ sec, ranging from 30°C to 55°C. To avoid plantar tissue damage, the upper cut-off limit was 20 sec. The left hind paw was evaluated three times, respecting a five minutes interval between then. Results were expressed in latency time average of the measurements (Hargreaves et al., 1988).

#### 2.3.3. Thermal cold allodynia

Cold allodynia was assessed by the acetone test, as previously described by (Kawashiri et al., 2011). Briefly, mice were placed in an acrylic chamber with a metal grid floor that allows free access to the animal’s paws. 30 min after the acclimation, 50 μl of acetone was sprayed onto the plantar skin and the nociceptive behavior (flinching and licking) time as recorded for five minutes.

#### 2.3.4. Formalin-induced nociceptive behavior

The formalin test was performed as previously described by (Hunskaar and Hole, 1987). After four weeks of diet intervention, control and zinc deficient groups received an intraplantar injection of formalin 2.5% (20 μL) on the right hind paw. The nociceptive behavior was recorded (licking time of the formalin-injected paw) for thirty minutes, first five minutes post-injection (neurogenic phase) and for fifteen minutes starting at the 15^th^ min post-injection (inflammatory phase).

#### 2.3.5. Carrageenan-induced hypernociception

Inflammatory pain was evaluated using carrageenan-induced mechanical hypernociception assay. After four weeks of diet intervention, mice were injected with 50 μl of 1% carrageenan solution in sterile saline (NaCl 0.9%) into the sub plantar surface of the right hind paws. The mechanical sensitivity was evaluated using calibrated von Frey filaments. Mice were placed individually in acrylic cages with wire grid floors and the withdrawal response was determined at 0, 1, 3, 6 and 24 h postchallenge.

### 2.4. Western Blot Analysis

Four weeks after diet intervention, the animals were euthanized and the lumbar dorsal root ganglions (DRGs) were collected. DRGs samples were homogenized in lysis buffer (1% Triton, 20 mM Tris pH 7.6, 1 mM EDTA, 150 mM NaCl, 1 mM NaF, 1 mM NaVO3) with protein inhibitor cocktail (Sigma®) followed by total protein quantification. Protein samples were resolved in SDS-polyacrylamide and transferred to a nitrocellulose membrane. The membrane was then blocked with TBS buffer (25mM Tris-HCl pH 7,6, 0,2M NaCl and Tween 20 0,15%) + 3% BSA for 1 hour, to reduce unspecific ligation, followed by incubation, with specific primary antibody - anti-SOD1 (Santa Cruz, RRID:AB_640874), anti-ATF-3 (Santa Cruz, RRID:AB_2258513), anti-GAP-43 (Santa Cruz, RRID:AB_640874) or anti-β-actin/actin (Santa Cruz, RRID:AB_2223228) - overnight at 4°C. After that, the anti-rabbit secondary antibody was incubated for 1 h and chemiluminescence was measured using a ChemiDoc XRS+ imaging system (Bio-Rad) after development in chemiluminescence reagent (Westar Nova 2011, CYANAGEN). Bands intensity was determined using ImageJ software (NIH, United States).

### 2.5. Plasma TNF quantification

To quantify the levels of TNF, animals blood were collected four weeks after the beginning of diet intervention by cardiac punction. Whole blood samples were centrifuged (10,000 rpm, 10 min at 4°C) and the plasma fraction collected to proceed the cytokine and protein measurements. TNF levels were determined using an enzyme-linked immunosorbent assay kit commercially available (R&D Systems®).

### 2.6. Metabolomics

#### 2.6.1. Metabolite extraction

Four weeks after diet intervention, the animals were euthanized and the livers were collected. Liver samples were kept in liquid nitrogen until use. For metabolite extraction, specimens were transferred to 2 mL microtubes and 1 mL of acetonitrile was added. Samples were mechanically disrupted with zirconia/silica beads (0.5 mm diameter) using a Mini-Beadbeater (1-min pulses, three times, with a 1-min rest on ice between pulses). The samples were centrifuged for 5 min at 16,000g and 4°C and the supernatants were carefully transferred to new 2 mL tubes, dried in a SpeedVac evaporator and stored at −80°C for further analysis.

#### 2.6.2. Mass spectrometry analysis

Electrospray ionization–mass spectrometry (ESI–MS) analysis was performed in a Traveling Wave Ion Mobility Mass Spectrometer (TWIM-MS, Synapt G1 HDMS, Waters, UK). Dried samples were solubilized (10 μL of 60% acetonitrile for each mg of initial liver sample), kept in an ultrasonic bath during 5 minutes, and centrifuged at 14,000 rpm and 6 °C during 30 minutes. Each sample was divided into two parts for analysis conditions in positive and negative mode. For analysis in positive mode was added formic acid at 0.1% and for negative mode ammonium hydroxide at 0.5%. The quality control analysis was performed with a mixture of all samples which were injected at the beginning and the end of the analysis. Mass calibration was performed with phosphoric acid solution in range from 50 to 2,000 m/z. Samples were injected at a rate of 10 μL min^-1^, data were acquired over the range from 100 to 1,000 m/z during 5 minutes in dynamic mode and scan time of 0.8 seconds. The positive ESI analysis was performed with a capillary voltage of 3.0 kV, sampling and extraction cone respectively at 50 V and 4.0 V. The settings for negative analysis were capillary voltage of 2.5 kV, sampling and extraction cone respectively at 50 V and 5.0 V. The nanoflow gas was set at 500 L h-1 at a desolvation temperature of 150 °C, and the source temperature set at 100 °C. All the acquisition of data was operated by Waters MassLynx v4.1 software.

Mass spectrometric data were processed using integrated MassLynx tools and MarkerLynx™ XS Application Manager from Waters (Milford, MA, USA). Firstly, raw data were centroided using the Accurate Mass Measure tool and then the resulting mass data was processed using MarkerLynxTM. The method parameters were as follows: analysis type = combined scan range, peak separation (Da) = 0.02 and marker intensity threshold (counts) = 2,000. Data were normalized to the total marker intensity and a two-dimensional data matrix (m/z versus peak intensity) was generated for each ionization mode and exported to a format amenable for further data analysis. Data from positive and negative modes were combined and Principal Component Analysis (PCA) was performed using the freely available software Multibase (http://www.numericaldynamics.com). To identify differences in metabolite composition between groups and to assign possible metabolite identities to m/z values showing at least a 2-fold difference in intensity between sets of samples we manually calculated the mean and standard error of mean (SEM) for each metabolite and data were queried against PATHOS – The metabolomics tool from Glasgow Polyomics (http://motif.gla.ac.uk/Pathos/index.html).

### 2.7. Data Analysis

Statistical analyses were performed using GraphPad Prism ver 7.01 (GraphPad Software). One-way ANOVA or *t*-test was used depending on the experiment. Bonferroni post-test was used in all analysis. A *p*-value of <0.05 was considered significant.

## 3. Results

### 3.1. Reduction of zinc intake since the weaning affects mice growth and body weight gain

Animals fed with control diet showed typical pattern of growth as indicated by both gain in body weight (F(4,64)=15.72; p<0.0001) (**Figure 1A**) and tibia length (t(8)=5.87; p=0.0004) (**Figure 1B**). In contrast, animals on zinc-deficient diet showed significantly restriction in growth that are also significant different from diet control animals (control diet = 23.2 ±1.4 g *versus* Zn-def diet = 14.9 ±1.1 g, 28 days after diet intervention; F(9,113)=16.16; p<0.0001) (**Figure 1A**). These data are in accordance with a priori knowledge about the effect of zinc restriction on animal growth, indicating the effectiveness of the zinc-restricted diet in achieving these expected outcomes. It is important, however, to highlight that zinc-deficient group also presented bowel malfunction, repetitive jumping behavior, hair loss and an increase of mortality (*data not shown*), confirming the impact of zinc deficiency to animal health development.

**Figure 1:**
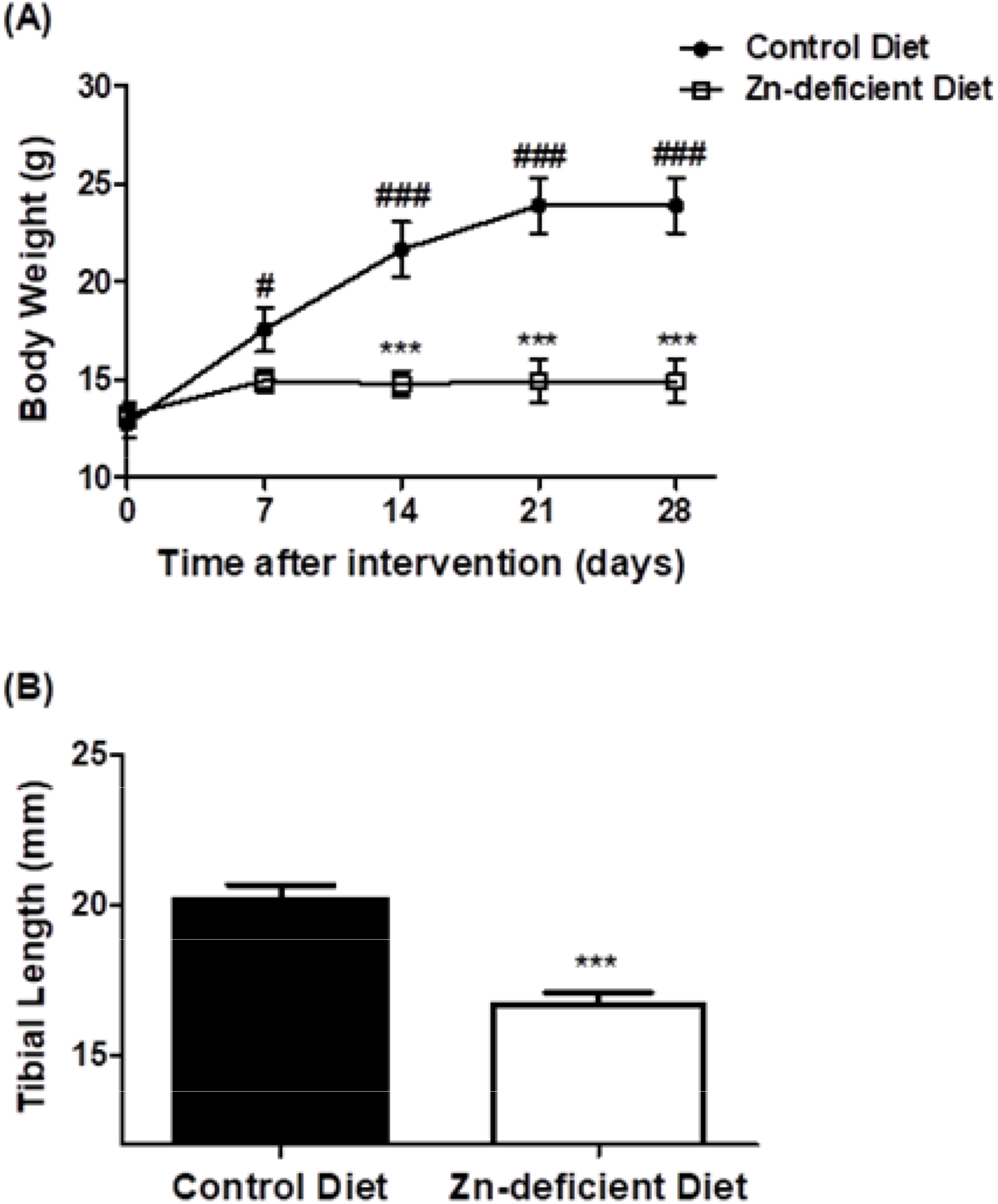
Effect of dietary zinc deficiency on mice body weight and growth. (A) Control (filled circle) and zinc-deficient (opened square) diet animals body weights were measured weekly. (B) Tibial length sizes were measured at week 4. Data are expressed as mean ± SEM, n=9-15 animals, One-way ANOVA followed by Bonferroni post-test (body weigth) and *t*-test (tibia length), ***p<0.001 vs control diet group, #p<0.05 and ###p<0.001 vs its baseline value.

### 3.2. Dietary zinc deprivation increases nociceptive behavior evoked by mechanical and thermal stimulus

To understand the impact of zinc-deficient diet on the basal nociceptive sensitivity, the response to mechanical and thermal stimulus were evaluated in mice (**Figure 2**). Basal mechanical sensitivity was measured weekly by the *von Frey* test (Chaplan et al., 1994). The response to thermal heat and cold stimulus was evaluated four weeks after the start of diet intervention by Hargreaves and acetone tests, respectively.

**Figure 2:**
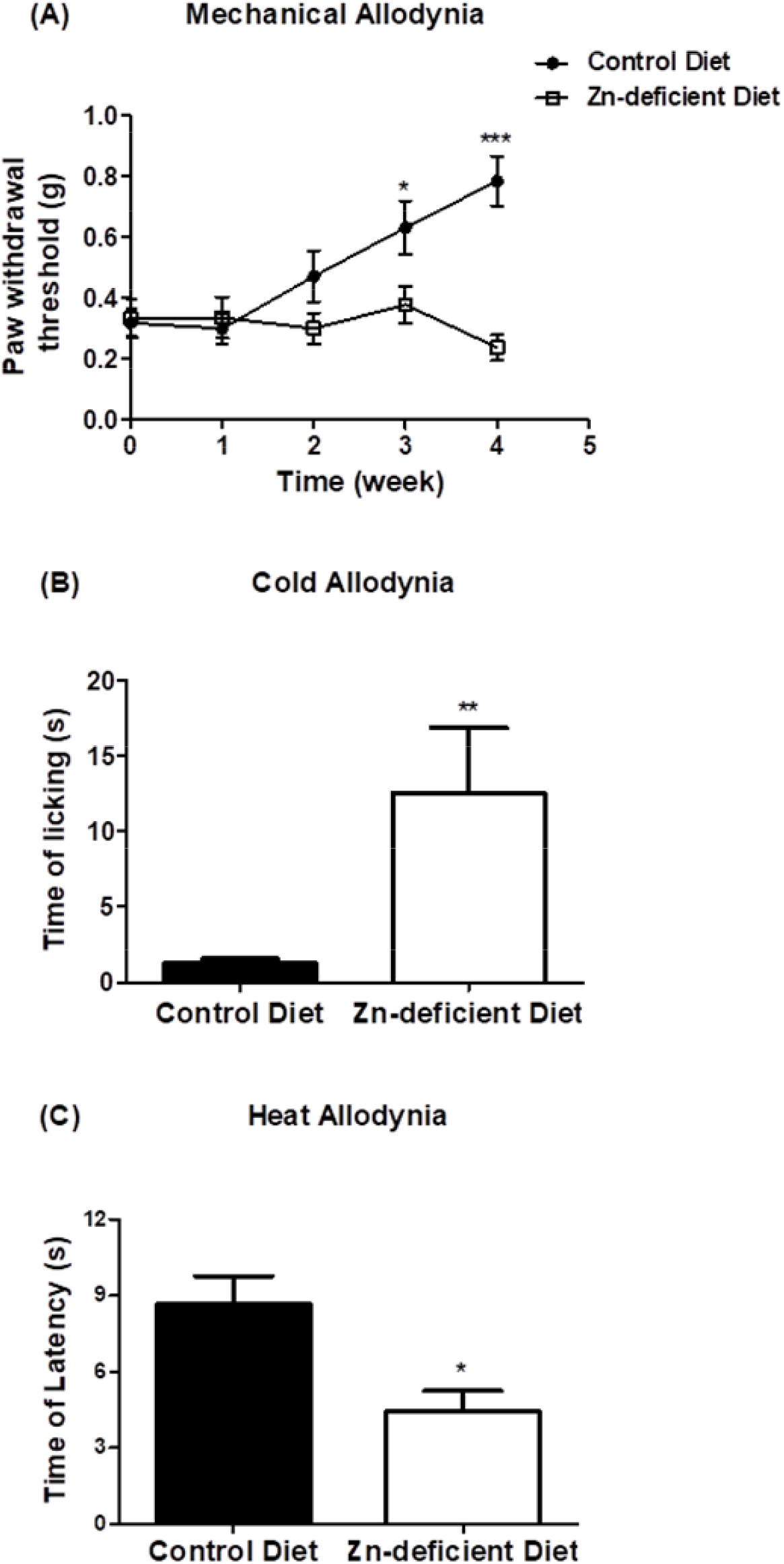
Effect of dietary zinc deficiency on mice mechanical and thermal allodynia. (A) Control (filled circle) and zinc-deficient (opened square) diet groups mechanical withdrawal threshold were measured weekly by *Von Frey* filaments. (B, C) Cold and heat sensitivity, respectively, were measured at week 4. Data are expressed as mean ± SEM of animals paw withdrawal threshold (g) (n=10-14), time of licking (s) (n=10-14) or time of latency (s) (n=4-5), One-way ANOVA followed by Bonferroni post-test (mechanical allodynia vs its baseline value) and *t*-test (thermal allodynia vs control diet group), *p<0.05, **<0.01 and ***p<0.001.

Young mice submitted to a control diet presented a low mechanical threshold that increased over the weeks along with growth (F(4,69)=8.12; p<0.0001) (**Figure 2A**), which is a normal pattern of animal development (McKelvey et al., 2015). In opposite, animals under zinc-deficient diet showed a dissimilar pattern with the mechanical threshold consistently remained low throughout the growth curve. After three weeks under dietary intervention, the control group showed a mechanical paw withdrawal threshold (PWT) of about 0.632±0.089g while the zinc-restricted group showed a PWT of about 0.379±0.061g (F(9,136)=6.59; p<0.0001). Mechanical allodynia is still observed to zinc deficient group at week four.

Mice submitted to a zinc deficient diet also presented a significant hypersensitivity to thermal stimulus after four weeks. The zinc-restricted group showed an increase in the nociceptive behavior evoked by cold stimulus (t(22)=3.135; p=0.0048) and a reduction of the time to respond to heat innocuous temperatures compared to control group (t(7)=2.93; p=0.022) (**Figure 2B** and **Figure 2C**). Together, these findings showed that zinc deficient diet promotes allodynia evoked by mechanical, cold and heat stimulus.

### 3.3. Zinc-deficient diet increases neurogenic pain but abolishes inflammatory pain behavior

Since the dietary zinc restriction resulted in disrupted basal sensitivity, we hypothesized whether animal’s hyperalgesic response could also be altered. To address this question we performed the formalin inflammatory test (Hunskaar and Hole, 1987).

The intraplantar administration of formalin induces two phases of pain behavior: neurogenic phase (0 to 5 minutes after formalin injection) and inflammatory phase (15 to 30 minutes after formalin injection). The first phase is characterized by activation of channels involved on pain behavior as TRPs and the second by the release of inflammatory mediators (prostaglandins, cytokines, etc.) that support the nociceptive response (Hunskaar and Hole, 1987).

Mice under zinc-deficient diet for four weeks showed an increase of the nociceptive behavior (licking time) in the neurogenic phase. The hypernociceptive response was about 50% higher in zinc-restricted mice than in regular diet group (t(14)=3.46; p=0.0038). However, in the inflammatory phase there was a significant reduction in the nociceptive response among animals with restricted zinc diet and control diet (t(14)=2.85; p=0.0128) (**Figure 3A**).

**Figure 3:**
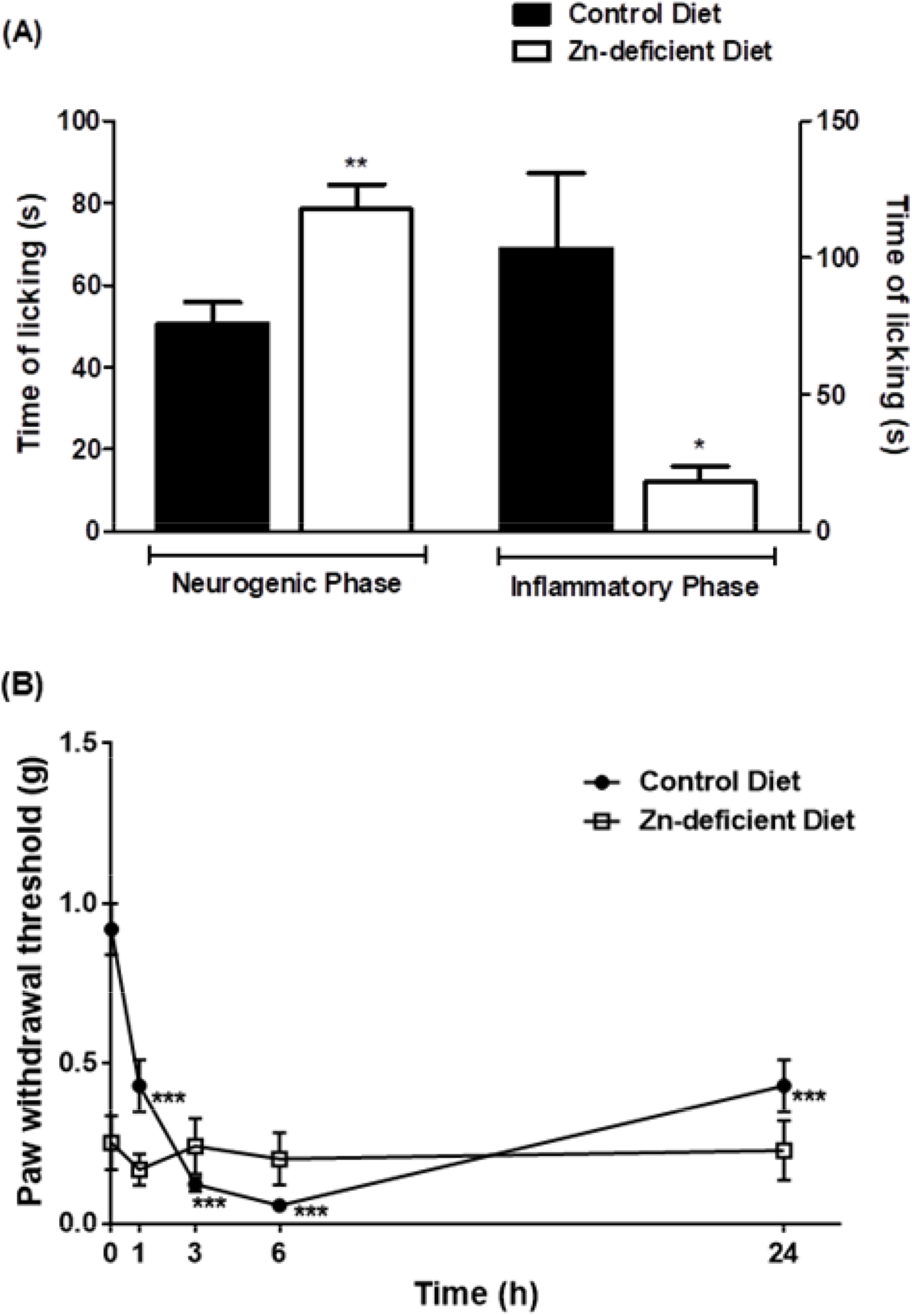
Effect of dietary zinc deficiency on formalin induced nociception (A) and mechanical hyperalgesia induced by carrageenan in mice (B). (A) Neurogenic and inflammatory phases of the formalin test (n=7-9). (B) Carrageenan induced hyperalgesia test (n=5-6). Time of licking (s) and paw mechanical withdrawal threshold (g) of control (filled bar/circle) and zinc-deficient (opened bar/square) diet groups were accessed at week 4. Data are expressed as mean ± SEM, *t*-test (licking time vs control diet group) and One-way ANOVA followed by Bonferroni post-test (paw withdrawal vs its baseline value), *p<0.05, **p<0.01 and ***p<0.001.

Considering that zinc restriction affected inflammatory pain in formalin test, we hypothesized if the reduction of zinc intake could also affect pain behavior after intraplantar injection of carrageenan, a classic inflammation inducer. As expected, animals that received control diet presented a significant reduction of mechanical PWT one hour after the inflammatory stimulus. The hyperalgesic behavior remains even twenty-four hours after carrageenan injection (F(4,20)=28.81; p<0.0001) (**Figure 3B**).

As already demonstrated (**Figure 2A**), zinc-deficient group shows a reduced basal mechanical sensitivity compared to animals that received regular amounts of zinc. This reduction was not modified by carrageenan administration (**Figure 3B**). PWT remained low, similar to its baseline and unchanged for twenty-four hours, showing that changes in sensitivity threshold promoted by dietary zinc deficiency were not affected by inflammation.

### 3.4. Reduction of zinc intake changes the expression of proteins related to glial satellite cell, neuronal activation and oxidative stress in DRG

In order to understand the mechanisms behind the hypernociceptive behavior showed by animals submitted to restriction in dietary zinc, we quantified the dorsal root ganglion (DRG) levels of proteins related to glial satellite cells (GFAP, *glial fibrillary acidic protein*), neuronal activation (ATF-3, *activating transcription factor 3*) and oxidative stress (SOD1, *superoxide dismutase 1*). DRGs were removed four weeks after diet intervention and submitted to western blot analysis.

A wide variety of neuronal stress conditions are able to induce a state of activation of glial satellite cells and neurons, which can be inferred by the evaluation of GFAP and ATF-3 expression, respectively (Galbavy et al., 2015; Malaspina et al., 2010). DRGs of animals under dietary zinc restriction presented a significant reduction of GFAP levels (t(6)=2.716; p=0.0348) (**Figure 4A**) accompanied by increased levels of ATF-3 (t(8)=6.107; p=0.0003) (**Figure 4B**) suggesting that neuron activation and damage occurs independently of satellite cells activation.

**Figure 4:**
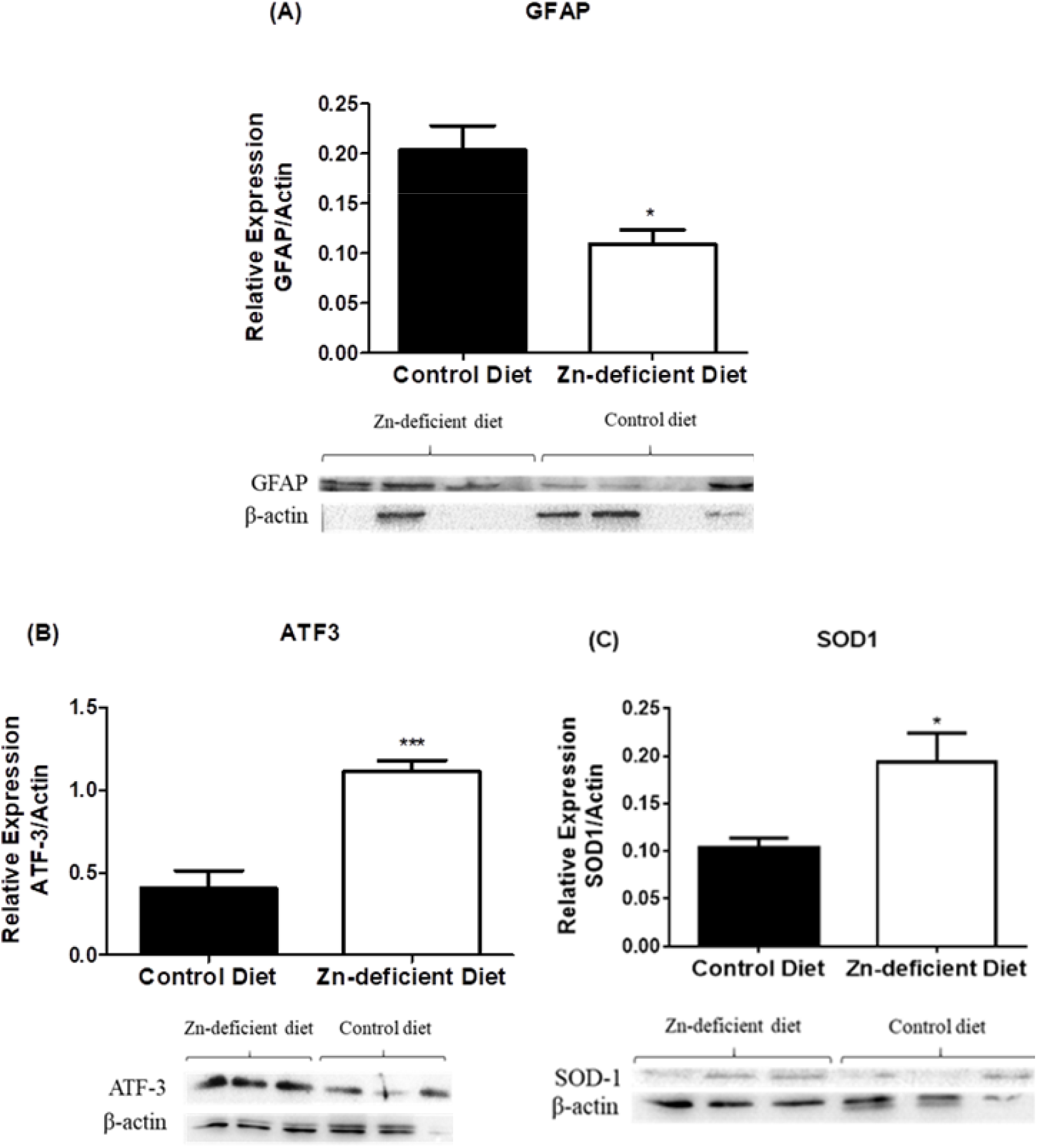
Zinc-deficient diet effects on glial satellite cells, neuron activation and oxidative stress. Lumbar DRGs were collected four weeks after diet and the levels of GFAP (A), ATF-3 (B) and SOD1 (C) were measured by western blot. Data are expressed as mean ± SEM of relative expression of GFAP, ATF-3 or SOD-1/actin, n=3-5 DRG, *t*-test, *p<0.05 and ***p<0.001 vs control diet group.

Aiming to understand if oxidative stress could be related with neuron activation and hypernociceptive disorders observed upon dietary zinc restriction, we also evaluate the levels of SOD1, an antioxidant enzyme that plays important role on oxidative stress and neuronal damage control. SOD1 levels were increased in the DRG of zinc-deficient mice compared to control group (t(4)=2.795; p=0.049) (**Figure 4C**).

### 3.5. Zinc-deficient diet decreases TNF plasma levels

In the present intervention model, zinc dietary intake reduction had a massive impact on inflammatory pain response, which prompted us to characterize if the downregulation of inflammatory process was taking place systemically. Since we observed the likeliness for increasing in neuronal stress with no effects in glial cells activation along with the absence of pain behavior in inflammatory conditions, we also evaluate the levels of the pro-inflammatory cytokine TNF.

Animals under zinc-deficient diet showed reduced plasma levels of TNF, compared to mice fed control diet (t(4)=7.574; p=0.0016) (**Figure 5**). This data corroborates the findings that inflammatory markers are downregulated in zinc-deficient group, providing evidences that nociceptive symptoms promoted by dietary zinc restriction in mice are unrelated to the increase in pro-inflammatory cytokines.

**Figure 5:**
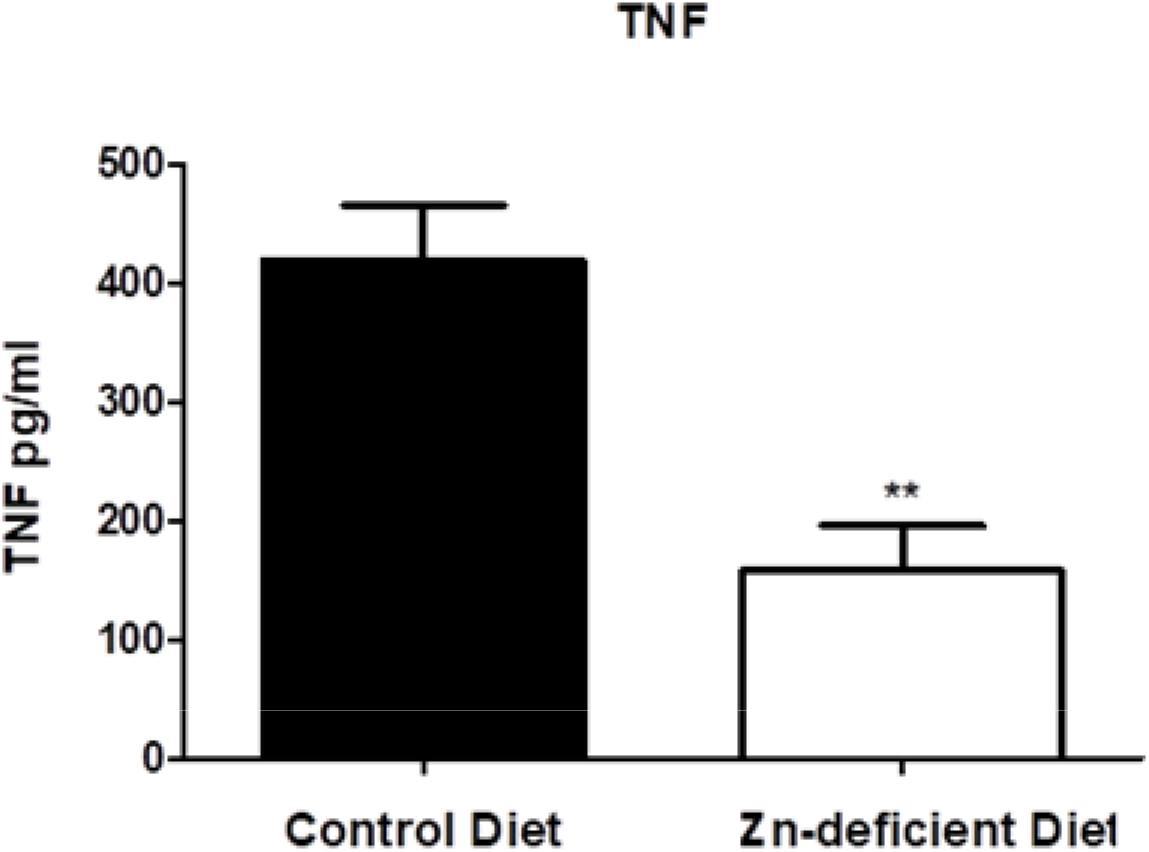
Zinc-deficient diet effect on TNF plasma levels. TNF plasma levels were measured four weeks after starting the diet. Data are expressed as mean ± SEM of TNF in pg/ml, n=3 animals, *t*-test, **p<0.01 vs control diet group.

### 3.6. Zinc-deficient diet downregulates arachidonic acid (AA) metabolism

To study the effects of zinc deficient diet on mice liver metabolic profile, ESI-QTOF-MS was used to detect and relatively quantify small metabolites in livers of animals fed with control or zinc-deficient diets. Such high-throughput analysis yielded a total of 3,021 different *m/z* (metabolic features) from both control and experimental groups which were detected from combined positive and negative ion modes (**Table 1**). A Principal Component Analysis (PCA) loading and score plots (**Supplementary Information S1A, S1B** and **S2**) illustrates the extensive differences in metabolic composition between livers from animals fed with control diet and zinc deficient diet.

**Table 1.**
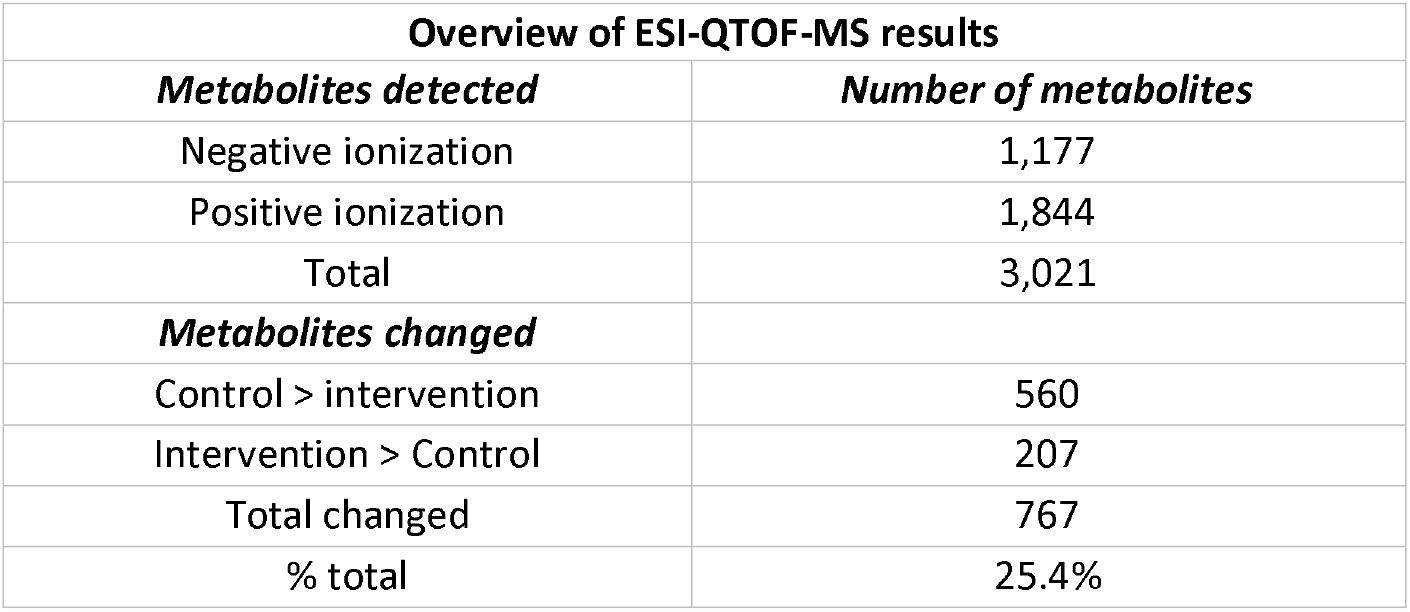
Liver metabolites quantification of control and zinc-deficient diet mice.

A more refined analysis was then carried out to determine the extent of the metabolic differences between animals subjected to control and zinc-deficient diets. In order to investigate which of the metabolites detected were present at different levels in those groups of samples, the average intensities of all metabolites were calculated and results from each of the sample groups (control and low zinc diet) were compared. Metabolites that showed changes of 2-fold or more were used for further analyses. Based on this analysis we found that 767 out of the total 3,021 metabolites were present at different levels and changed by the diet when comparing samples from different diets (**Table 1**). This represents 25.4% of all detected m/z, providing evidences that an extensive metabolic shift occurs on animals fed with zinc restriction.

The metabolic pathways most significantly disturbed in livers and that differ between the two groups were illustrated in **Figure 6**. Although many metabolic pathways were affected, our data suggest that the arachidonic acid (AA) metabolism is markedly modulated due to zinc-deficient diet, presenting lower levels of many of its metabolites when compared to the control. Two putative metabolites of AA pathway showed increased levels in that experimental group (TXB2 and 6-Keto PGF1α) and only one showed no difference between the two groups (LTD4), as depicted in the AA metabolic pathway (**Figure 7**). Although the AA itself was not detected experimentally, several m/z corresponding to an array of potential AA derivatives were detected in lower levels (two-fold or higher) in mice livers with zinc restriction. The relatively high number of metabolites of AA pathway that were identified by our exploratory method suggests that this is indeed an important component of immune response disruption promoted by dietary zinc restriction, indicating that this metabolic route generates a diverse class of bioactive lipid mediators involved in inflammatory and pain processes.

**Figure 6:**
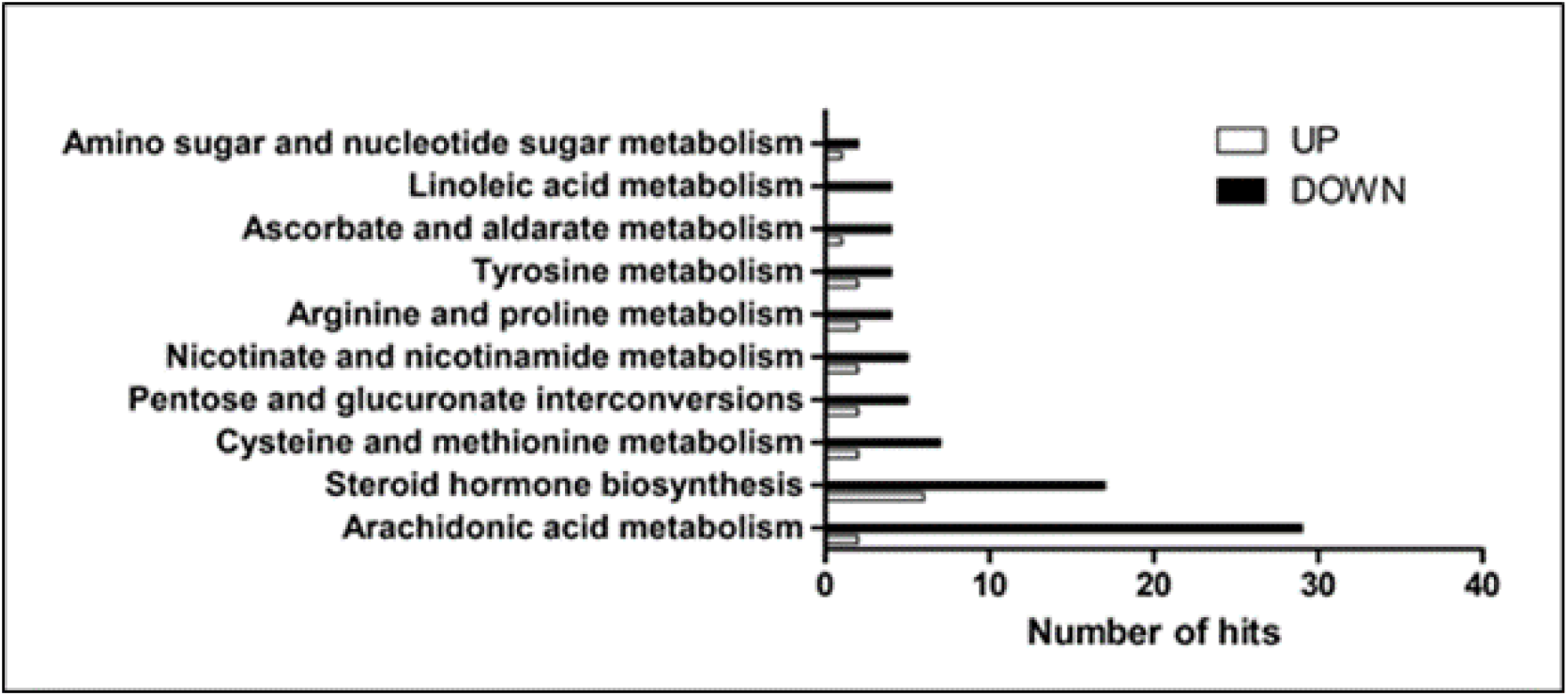
Metabolomics analysis of liver from control and zinc-deficient mice. Total number of hits from metabolomic pathways altered upon intervention with zinc-deficient diet. Ions detected from electrospray analysis in both positive and negative modes were searched in the KEGG database and represented here for those that increased (open) or decreased (closed) compared to animals under control diet.

**Figure 7:**
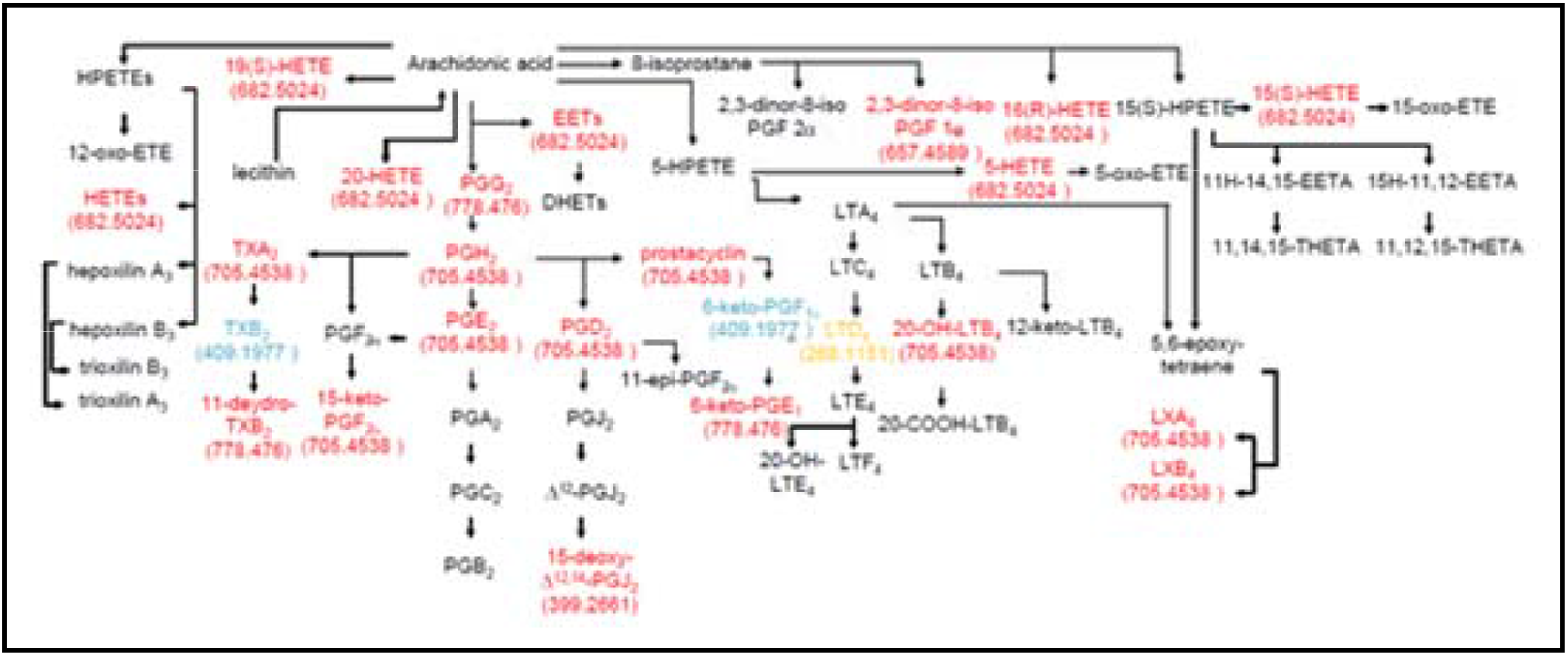
Effect of dietary zinc deficiency on the arachidonic acid metabolism in mice. Schematic overview of the arachidonic acid metabolism pathway is show, depicting the metabolites (red) increased or (black) unaltered in relative abundance upon dietary zinc restriction compared to animals fed with control diet. Numbers in parentheses represents the true *m/z* detected (for the M:H) ion.

## 4. Discussion

The present work shows the effect of a zinc deficient dietary on the nociceptive, inflammatory and behaviors responses in mice and its mechanism of action.

Zinc is recognized as a fundamental ion for proteins and enzymes, acting as a catalytic, co-catalytic factor or as a structural element (Bert I. Vallee; Falchuk, 1993). In recent years, the role of zinc in the development and functioning of the nervous system has been highlighted. Several studies have shown zinc as an essential micronutrient for the development of memory and learning processes (Frederickson et al., 2005; Sensi et al., 2011).

Zinc levels play important roles in painful and neurodegenerative conditions, as spinal cord injuries, being able to modulate the spinal cord signaling of pain (Ma and Zhao, 2001; Su et al., 2012). Despite to the growing evidences of the zinc participation in pain modulation, the knowledge about the consequences of zinc dyshomeostasis to pain disorders and its mechanisms are still unclear. Here in we showed for the first time the impact of dietary zinc intake reduction (one-third less than recommended AIN93 levels) to mice development nociceptive disorders with unanticipated decrease in inflammation pattern and metabolism.

Dietary zinc restriction, since the weaning, increased nociceptive behavior evoked by thermal and mechanical stimulus. These data corroborate previous findings on zinc modulation of pain evoked by different stimulus (Liu et al., 1999; Nozaki et al., 2011). Liu et al (1999) (Liu et al., 1999), demonstrated that zinc ions, delivered by intraperitoneal, intraplantar or intrathecal routes to rats with neuropathic pain, exert anti-nociceptive effects over thermal heat hyperalgesia. Furthermore, it has been showed that reduction of spinal zinc levels, by selective chelators, or of NMDA receptors zinc affinity, by mutation, promotes mechanical and thermal heat hypersensitivity (Larson and Kitto, 1997; Nozaki et al., 2011), reinforcing the relevance of zinc signaling to heat hyperalgesic responses.

Disruption of zinc homeostasis also increases the allodynic behavior evoked by cold temperatures in mice. Cold allodynia is a common symptom of different pain conditions, such as inflammatory and neuropathic, and several factors has been described as responsible for its modulation (Greenspan et al., 2004; Knowlton and Mckemy, 2011). The transient receptor potential (TRP) channels, specially TRPM8 and TRPA1, are well stablished as molecular sensors through cold stimulus, as well as the neurotrophic factors NGF (nerve growth factor) and GFL (glial cell line-derived neurotrophic factor family ligand) that are involved in cold hypersensitivity (Caspani et al., 2009; Lippoldt et al., 2016). The capacity of zinc to act in ion channels modulation together with its ability to regulate the central and peripheral levels of NGF, can in part explain the evoked cold allodynia symptoms presented by zinc deficient mice (Chowanadisai et al., 2005; Feske et al., 2015; Safieh-Garabedian et al., 1996).

Zinc deficiency leads an increase of licking time of the neurogenic phase of the formalin test, whereas it almost abolishes the pain behavior of the inflammatory phase. The painful behavior of neurogenic phase is the result of nociceptors activation by the chemical agent followed by a massive contribution of central sensitization (Lebrun et al., 2000). Zinc is an endogenous regulator of excitatory neurotransmission, being described as an allosteric modulator of NMDA receptors at spinal cord and able to attenuate C fiber-evoked potentials (Ma and Zhao, 2001; Nozaki et al., 2011). These evidences suggest that pro-nociceptive effects observed at neurogenic phase of formalin test, as well as the other painful behaviors observed in this work for zinc-deficient mice, are due to the loss of physiological control of pain by endogenous zinc.

The painful behavior of the inflammatory phase of formalin test happens as result of the mobilization of wide array of signaling molecules including neurotransmitters, peptides, eicosanoids and related lipids (prostaglandins, thromboxane, leukotrienes, endocannabinoids), neurotrophins, cytokines and chemokines (Basbaum et al., 2009). Immune cells are pivotal to the establishment of this “inflammatory soup” releasing and responding to several pro-inflammatory and pro-algesic mediators. Zinc is well recognized as an important factor to immune response regulation. Zinc deficiency has been associated with decrease of macrophages and neutrophils chemotaxis, phagocytosis and oxidative stress. Moreover, zinc levels seems to be important for the release of pro-inflammatory cytokines by macrophages (Haase et al., 2008; Ryu et al., 2011). In this sense, our findings suggest that reduced hypernociceptive behavior (inflammatory phase of formalin test) of zinc deficient group happens because the immune system response of these animals is compromised.

To better understand the effects of zinc intake reduction to the inflammatory pain we evaluate mice mechanical hypersensitivity evoked by carrageenan, a classical flogistic agent. In agreement with the formalin test findings, zinc deficient group presented a higher paw withdrawal threshold when compared to control group, confirming the lack of inflammatory response in this condition.

We also observed a downregulation of GFAP, an important astrocyte activation marker, in DRG of mice submitted to reduced levels of zinc, different of what is usually described in classical chronic pain conditions (Galbavy et al., 2015). On the other hand, zinc restriction promoted an increase of ATF-3, an well-known transcription factor upregulated during neuronal activation (Malaspina et al., 2010). In this sense, the increased ATF-3 level in DRG of zinc deficient mice shows a neuronal activation despite of glial cells downregulation. The reduction of serum TNF levels reinforced the idea of immune response disruption promoted by zinc deprivation. It also suggests that nociceptive symptoms observed are not dependent of peripheral TNF signaling. In consonance with neuronal distress and oxidative stress, characteristics linked to chronic pain, an important oxidative stress biomarker, SOD1, is also upregulated in DRG. Moreover, these findings reinforce the idea of immune response and oxidative stress disruption of sensory neurons during zinc deficiency conditions.

The impact of zinc deficiency on lipid metabolism has been demonstrated in different experimental models using rodents(Bettger et al., 1979; Johanning and Dell, 1989; Weigand and Egenolf, 2017). According to previous studies low zinc diets submitted animals to inflammatory conditions, such as dermal lesions, compatibles to fatty acids decrease (Bettger et al., 1979). Also, it has been shown that Zn-deficient diet reduces liver levels of arachdonic acid (Dieck et al., 2005). The impact of liver arachidonic acid reduction could be expanded to plasma and tissues surrounding nociceptors, affecting production of prostanoids which sensitizes these cells. In the present study, through a metabolomics analysis, we showed that zinc intake reduction since weaning was able to downregulate several metabolites of the acid arachidonic metabolism pathway that corroborate and explain the lack or reduced inflammatory pain behavior.

## 5. Conclusions

Zinc deficiency promoted by reduction of dietary intake since weaning induces nociceptive disorders that are independent of some classical pro-inflammatory and pro-algesic mediators. Thus, our findings reinforce the relevance of endogenous zinc levels balance to maintenance of sensory transmission homeostasis, and revels zinc deficiency as a potential risk factor to the onset of chronic pain disorders as painful neuropathies.

In addition to the previously described analgesic effect of Zn, we provide new evidence on the role of Zn on pain development, highlighting the clinical relevance of a dietary zinc supplementation.

## Supporting information

Supportng Material

## ACKNOWLEDGMENTS

The authors thank CAPES (BR), CNPq (BR), FAPERJ (BR) and INMETRO (BR) for financial support and fellowships.

## DATA AVAILABILITY

The datasets generated and/or analyzed during the current study are available from the corresponding author on reasonable request.

