## Supplementary material for "Dietary zinc restriction induces nociceptive pain with reduced inflammation in mice": Supportng Material

**SUPPORTING INFORMATION**

**S1. Comparison of the relative levels of liver metabolites identified in positive mode from control and zinc-restricted mice.** A) arachidonic acid pathway; B) steroid hormone biosynthesis. The metabolites listed correspond to those most probable to the identified *m/z*. Numbers are normalized to 100 % (compared to control) and numbers between parenthesis correspond to standard deviation.

A)

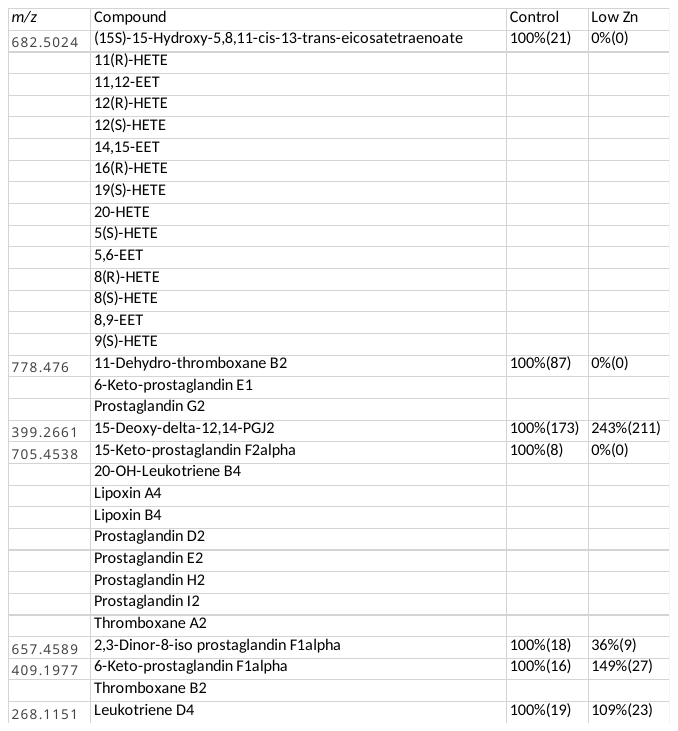

B)

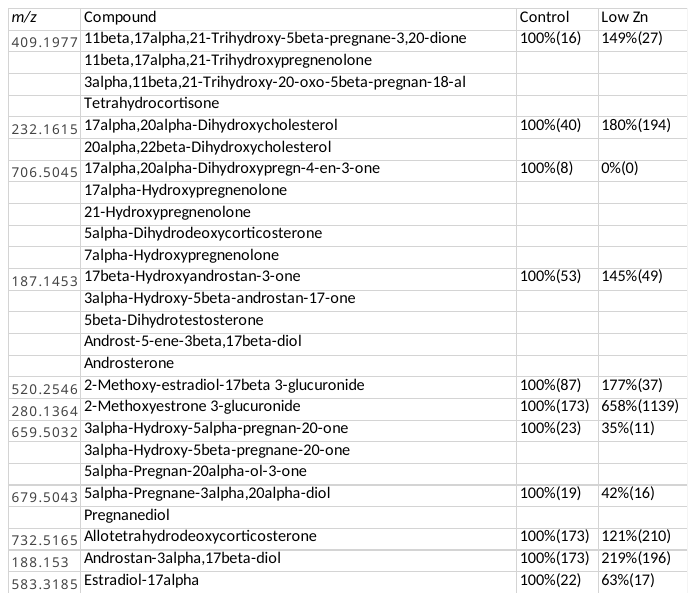

**S2. Principal Component Analysis of the Metabolomics Data.** Raw ESI-MS from both positive and negative ionization modes were combined and subjected to PCA analysis. The graphic representation corresponds to the control an intervention groups, and a combined sample from both groups acting as a monitor for data collection quality. Details in the *Material and Methods* section.

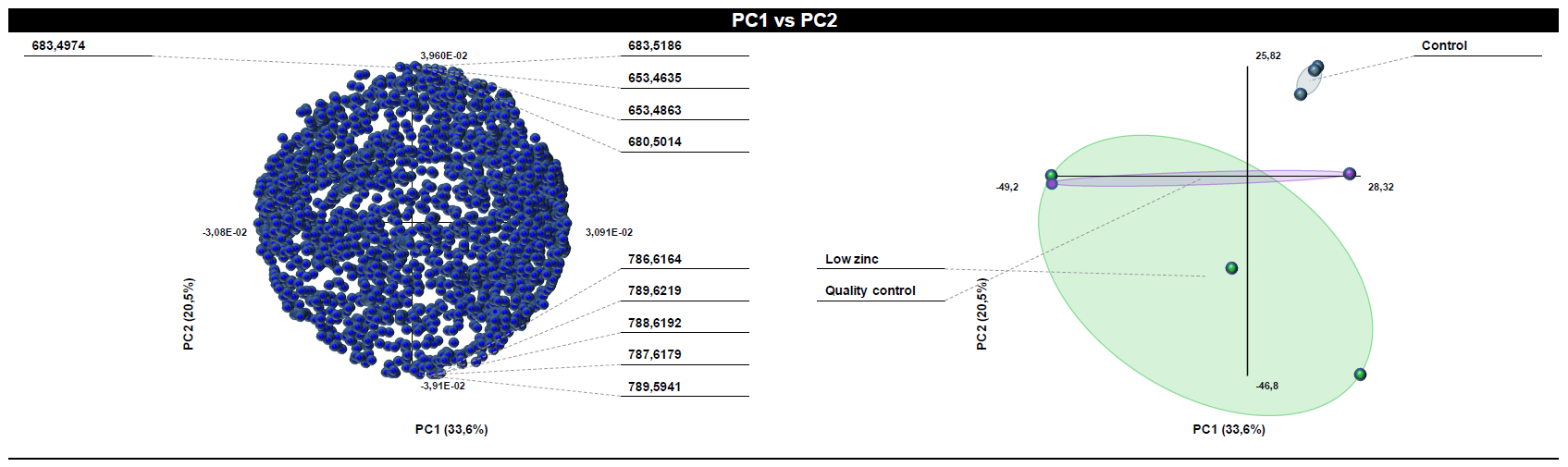
